# Lipid rafts increase to facilitate ectoderm lineage specification of differentiating embryonic stem cells

**DOI:** 10.1101/520106

**Authors:** Chen Xu, Bo Cao, Ying-dong Huo, Gang Niu, Michael Q Zhang, Zi-lin Mai, Xi-bin Lu, Han-ben Niu, Dan-ni Chen, Yan-xiang Ni

## Abstract

Lipid rafts are packed nanoscopic domains on plasma membrane and essential signalling platforms for transducing extracellular stimuli into cellular responses. Although depletion of raft component glycoshpingolipids causes abnormality particularly in ectoderm layer formation, it remains unclear whether rafts play a role in lineage determination, a critical but less-known stage in lineage commitment. Here, inducing mouse embryonic stem cell (mESC) differentiation with retinoic acid (RA), we observed lipid rafts increased since early stage, especially in ectoderm-like cells. Stochastic optical reconstruction microscopy characterized at super-resolution the distinct raft features in mESCs and the derived differentiated cells. Furthermore, RA-induced commitment of ectoderm-like cells was significantly diminished not only by genetic ablation of rafts but by applying inhibitor for glycosphingolipids or cholesterol at early differentiation stages. Meanwhile, raft inhibition delayed RA-induced pluripotency exit, an early step required for differentiation. Therefore, lipid rafts increase and facilitate ectoderm lineage specification as well as pluripotency exit during mESC differentiation.

## Introduction

Lipid rafts are compacted nanoscopic domains on plasma membrane and composed of saturated lipids, like glycospingolipids and cholesterol [1]. As such, rafts have been known to function as signalling platforms in transducing various extracellular stimuli into cellular responses [1-3]. Due to the compacted organization and nanoscopic sizes, rafts have so far failed to be well characterized by conventional techniques, such as confocal imaging and biochemistry assays. This limits the further insight into rafts’ new roles in a diversity of cellular events [2], although rafts have been implicated to exist in distinct cell types, including mouse embryonic stem cells (mESCs) [4, 5]. Deletion of glycosphingolipids through *ugcg* knockout caused abnormality particularly in ectodermal layer formation at E7.5 while having little impact before blastocyst stage [6]. These results implicate an essential role of glycosphingolipids/rafts in regulating proper lineage commitment(s) rather than self-renewal. Therefore, it will be interesting to explore the potential role(s) of lipid rafts in lineage commitment, particularly with the use of newly technique stochastic optical reconstruction microscopy (STORM) [7-9] that provides 10-fold better spatial resolution for resolving nanoscopic structures like rafts [10-12].

Lineage commitment has been studied *in vitro* in mESCs that are characterized by self-renewal and pluripotency [13]. mESC is capable of being committed into various specific cell types, each of which is contributed by coordination of multiple underlying gene regulatory networks (GRN). Balanced by pluripotency factors and various kinds of lineage factors, mESC remains at pluripotent state and will exit from pluripotency and go towards establishing a certain cell identity in response to lineage specifiers [14]. Intracellular signalling molecules, such as mitogen-activated protein kinase (MAPK), facilitates mESCs to exit from pluripotent state[15, 16]. Extracellular factors, such bone morphogenetic proteins (BMP) and fibroblast growth factor (FGF), influence cell identity establishment via functioning with their receptors on plasma membrane [17]. Although rafts have been known to mediate various receptor signalling pathways, it remains to be clarified if rafts function as signalling platform to contribute to pluripotency exit and lineage specification that have been studied relatively less. All-trans retinoid acid (RA) is important in embryonic differentiation *in vivo* [18] and has been widely applied in inducing mESC differentiation *in vitro* [19-22]. When added to mESCs, RA binds to its nuclear retinoic acid receptor (RAR) and regulate gene expression profiles, leading to pluripotency exit and multiple lineage commitments [20, 21]. Therefore, it will be interesting to investigate lipid rafts’ potential role in lineage specification via exposing RA to mESCs, the well-established *in vitro* differentiation model.

In this study, after exposing RA to monolayer mESC cultures, we found that cell-surface rafts increased since the 12 h time point, especially in ectoderm-like cells. We also adopted the emerging attractive technique STORM and characterized at super-resolution distinct nanoscopic features of rafts on the plasma membrane of mESCs and RA-induced differentiated cells. Furthermore, RA-induced commitment of ectoderm-like cells were significantly diminished not only by CRISPR/Cas9-mediated *ugcg* ablation, which resulted in raft depletion, but by inhibiting glycosphingolipids or cholesterol during the early differentiation. Meanwhile, raft inhibition delayed RA-induced exit of pluripotency, an early step required for lineage commitment. Collectively, lipid rafts increase and play a role in facilitating specification of ectoderm-like cells as well as pluripotency exit during RA-induced differentiation.

## Result

### Lipid rafts in mESC increase during RA-induced differentiation, particularly in ectoderm-like cells

To investigate the potential role of lipid rafts in lineage commitment(s), we applied a widely-used mESC J1, which was derived from inner cell mass (ICM) and observed to exhibit cell-surface lipid rafts by laser scanning confocal microscopy [23]. J1 cells cultured in monolayer were induced towards various lineages via exposing to RA [24, 25], an essential cue for lineage commitments *in vivo* and *in vitro* [18, 22]. After 96 h RA exposure, the cells demonstrated obvious morphological change from round colonies to flat and strongly adherent cells. To provide a portrait of lipid rafts on plasma membrane of mESCs or their differentiated cells, we labelled live cells with Alexa Fluor 647-conjugated cholera toxin B subunit (CTB AF647) that binds to raft component ganglioside GM1and has been widely used to label rafts [26-28]. For raft labelling, 18 °C was adopted to avoid the possible endocytosis or membrane domain aggregation that occurs at higher or lower assay temperature, respectively [28-31]. After 96 h RA exposure, cells were analysed by immuno-fluorescence staining of the key pluripotency factor Oct4 that decreased to a barely detectable level in differentiated cells (**Figure 1a,** leftmost). At the same time, we detected a dramatic enhancement of CTB binding in differentiated cells, compared to control J1 (**Figure 1a,** left), which was further confirmed by the quantitative results from flow cytometry analysis (**Figure 1b**). Confocal imaging revealed remarkably raft enhancement in RA-induced differentiated cells, regardless of different labelling temperatures (12°C, 18°C, and 24°C), and that the 18°C appeared optimal for raft labelling (**Supplemental Figure 1**). These findings indicated that lipid rafts exist in mESC and increase after RA exposure.

**Figure 1.**
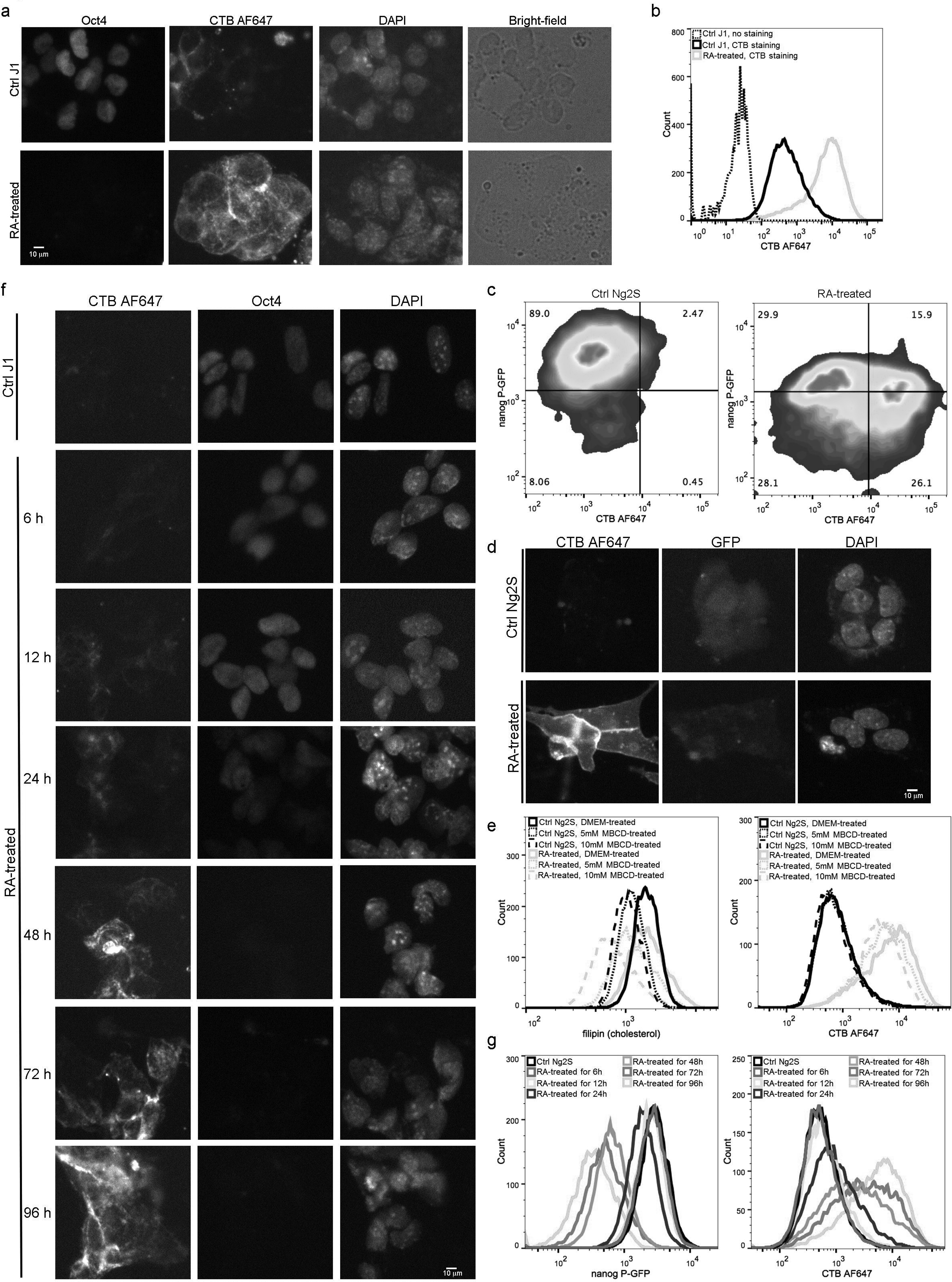
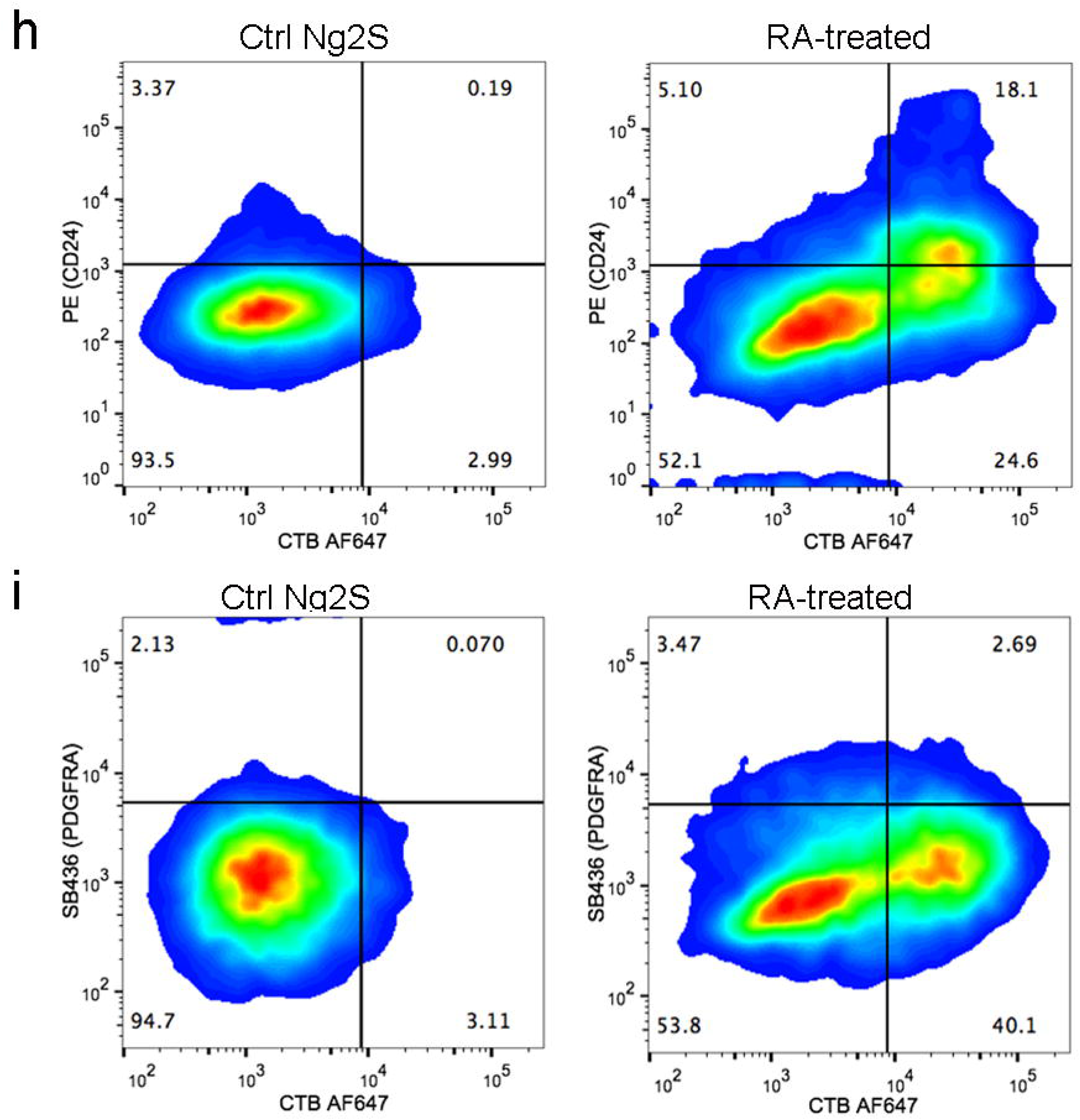
RA exposure to mESCs results in cell-surface raft increases, particularly in ectoderm-like cells. **a**, Immunofluorescence microscopy of RA-treated or control J1 stained with CTB AF647 and anti-Oct4 antibody. J1 cells were cultured with 1 μM RA or DMSO (Ctrl) for 96 h and labelled with CTB AF647. The fixed cells were then stained with anti-Oct4 antibody, followed by AF488 conjugated secondary antibody, and imaged by Nikon-Ti. Scale bars are 10 μm. **b**, Flow analysis of CTB labelling for J1 treated with RA (green) or control DMSO (black) for 96 h. The black dotted line represents negative control group without CTB labelling. **c**, Flow cytometry measurement of CTB labelling and Nanog P-GFP for Ng2S treated with RA or DMSO (Ctrl) for 96 h. The pseudocolor plots represent the relative density of cells. **d**, Immunofluorescence analysis of CTB labelling after cell fixation. Ng2S cells were cultured with 1 μM RA or control DMSO for 96 h. After fixation with 4% PFA, cells were labelled with CTB AF647. CTB labelling and nanog P-GFP were observed by Nikon-Ti using 641 nm and 488 nm, respectively. Scale bars are 10 μm. **e**, Flow cytometry measurement of filipin staining and CTB labelling for Ng2S with MBCD treatment. Ng2S cells cultured with 1 μM RA (green) or control DMSO (black) for 96 h were subsequently pulsed with 5 mM MBCD (dotted line), 10 mM MBCD (dashed line) or DMEM (solid line) for 30 min at 37°C. Then cells were labelled with CTB AF647 and fixed, followed by filipin staining. **f,g**, Dynamics of raft levels in mESCs treated with RA or control for a time course. The mESCs were exposed to 1 μM RA or control DMSO for 6 h, 12 h, 24 h, 36 h, 48 h, 72 h or 96 h and labelled with CTB AF647 before fixation. The J1 group samples were then stained with anti-Oct4 antibody, followed by AF488 conjugated secondary antibody, and imaged using Nikon-Ti (**f**). Scale bars are 10 μm. Ng2S group samples were subjected to **f**low analysis for quantifying of nanog P-GFP expression and CTB labeling (**g**). **h**, Flow cytometry measurement of CTB labelling, CD24 and PDGFRA expression of Ng2S treated with or without RA. Ng2S cells cultured with 1 μM RA or control DMSO for 96 h were labelled with CTB AF647 before fixation. Cell samples were co-stained with anti-CD24 and anti-PDGFRA antibodies. The pseudocolor plots represent the relative density of cells. Ctrl, Control; Exp, Experiment

To test if this observed alteration is constricted to J1, we applied another two mESC lines, the wide-type Cj9 and the genetically engineered Ng2S cell that expresses GFP reporter under the control of nanog promoter [32]. Consistently, profound increase of cell-surface rafts appeared in RA-induced differentiated cells, the differentiation state of which was confirmed by flow analysis of GFP expression in Ng2S (**Figure 1c**) or Oct4 staining in Cj9 (**Supplemental Figure 2**). Among RA-treated cells (right panel), the most differentiated subpopulation with decreased GFP level exhibited remarkable increase of CTB binding (**Figure 1c**), implicating a potential correlation between raft increase and cell differentiation. The observed increase was not a CTB-mediated tetrameric artefact caused by live cell labelling, as similar increase was observed when CTB staining was performed after fixation (**Figure 1d**). Since lipid rafts are maintained by the tight associations among lipid components, glycosphingolipids and cholesterol [33, 34], we analysed if RA-induced raft increase was abolished by cholesterol depletion with the use of methyl-β-cyclodextrin (MBCD) [35, 36]. As shown in left panel of Figure 1e, cellular cholesterol was efficiently depleted through transient MBCD treatment. MBCD treatment at two different concentrations significantly diminished the increase of CTB binding in RA-induced differentiated cells (**Figure 1e**, right, green), confirming that the increased CTB signal in differentiated cells reflects an enhanced abundance of rafts. Meanwhile, little difference was detected in control mESCs treated with or without MBCD (**Figure 1e**, right, black), which is probably due to the low basal level of cell-surface rafts in the control mESCs Taken together, these results demonstrated a profound difference in raft abundance between RA-induced differentiated and control mESCs.

Next, we examined the dynamics of cell-surface raft levels by treating mESCs with RA in a time course. In J1 cells, Oct4 expression was found dramatically reduced after RA exposure for 36 h-48 h (**Figure 1f**, middle column), which is known as exit from pluripotency with strong down-regulation of pluripotency markers [37]. Meanwhile, rafts increase appeared since the 12 h time point and augmented profoundly with the increase of RA exposure time (**Figure 1f**, left column). Flow analysis of RA-treated or control Ng2S provided quantitative evidence that raft increase was undetectable until 12 h after RA treatment (light blue line), preceding the exit from pluripotency (**Figure 1g**). Therefore, rafts increase from the very early differentiation stage.

Since RA treatment of monolayer mESCs led to the differentiation into ectoderm and extraembryonic endoderm (XEN)-like lineages [37], we explored the potential relationship of raft increase and the committed lineage by co-staining rafts with CD24 and PDGFRA, markers for ectoderm-like and XEN-like cells, respectively. In CD24-positive ectoderm-like cells, a main subpopulation demonstrated increased raft level (**Figure 1h**), indicating a possible relationship between ectoderm-like cell commitment and increase of raft level. Meanwhile, the PDGFRA-positive XEN-like cells exhibited no preferential to higher raft level (**Figure 1i**). These results suggested that lipid rafts might plays a role in mESC commitment to ectoderm- rather than XEN-like cells.

### Distinct nanoscopic features of cell-surface rafts in mESCs and RA-induced differentiated cells

Although conventional fluorescence microscopy and flow cytometry demonstrated that the abundance of cell-surface rafts increased during RA-induced differentiation (**Figure 1**), they failed to reveal the detailed features of these nanoscopic structures. In this regard, we sought to STORM that provides better spatial resolution, e.g lateral resolution of 22 nm in our lab [38]. STORM breaks optical diffraction limit via stochastically activating switchable fluorophores like Alexa Fluor 647 (AF647) at separate times, and the final super-resolution images are reconstructed with the measured positions of these individual fluorophores [7-9]. Here, bright field and conventional fluorescence images of J1 cells labelled with CTB AF647 were captured first (**Figure 2a**, left, upper). Then, sequential images were collected for super-resolution image reconstruction. Unlike the obscure structures imaged under conventional fluorescence microscopy (left, upper), discrete nanoscopic domains were uncovered to distribute on plasma membrane in STORM images, mostly with diameters no more than 200 nm (rightmost, upper, **Figure 2a**). In the control cell samples stained with AF647-conjugated IgG, no AF647-positive domains were detected (**Figure 2a**, rightmost, bottom), excluding that the identified nanoscopic structures were due to non-specific association of AF647 with the membrane. Since CTB mainly recognizes lipid raft marker, ganglioside GM1, these results confirmed rafts’ existence on cellular plasma membrane [10-12].

**Figure 2.**
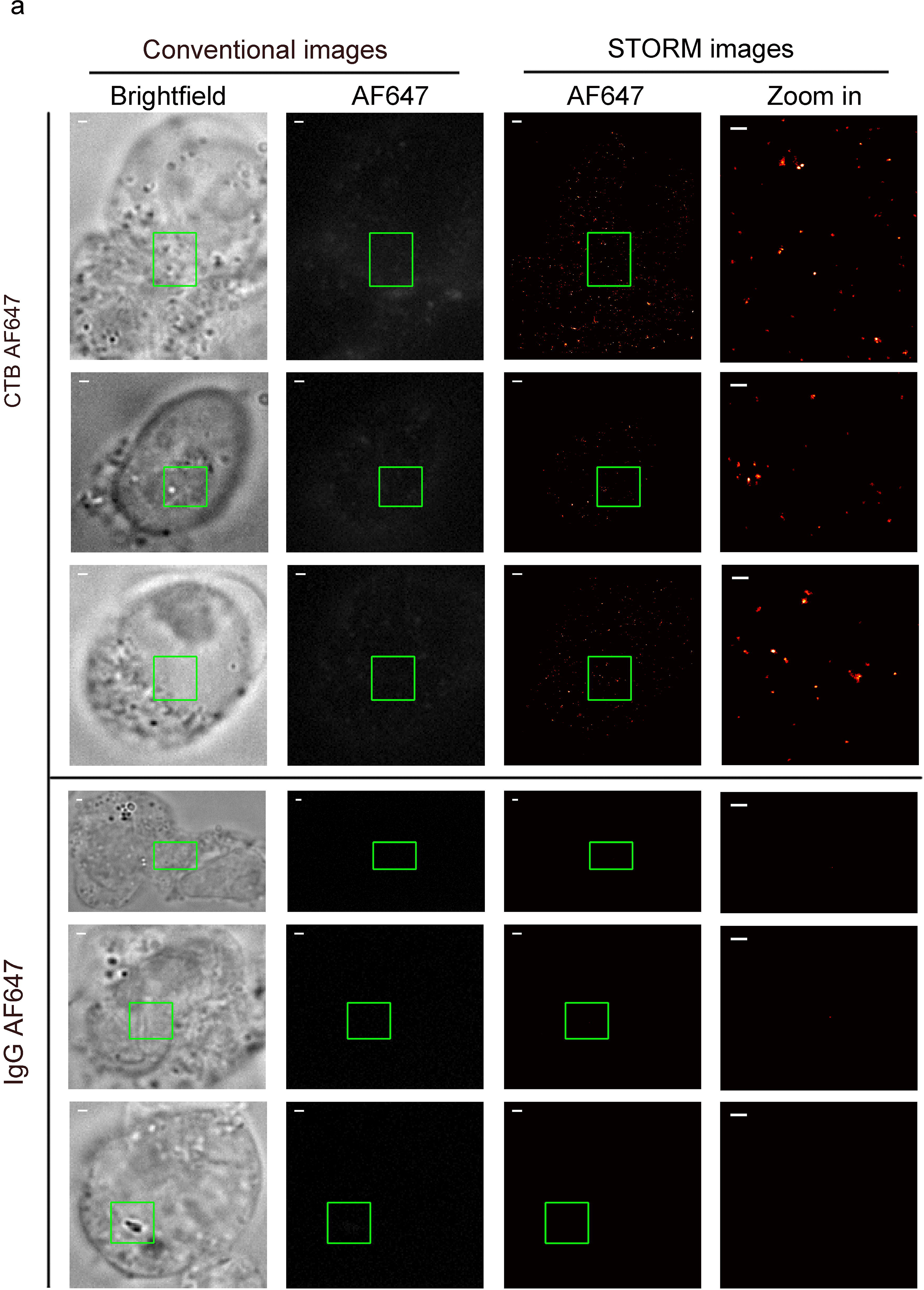
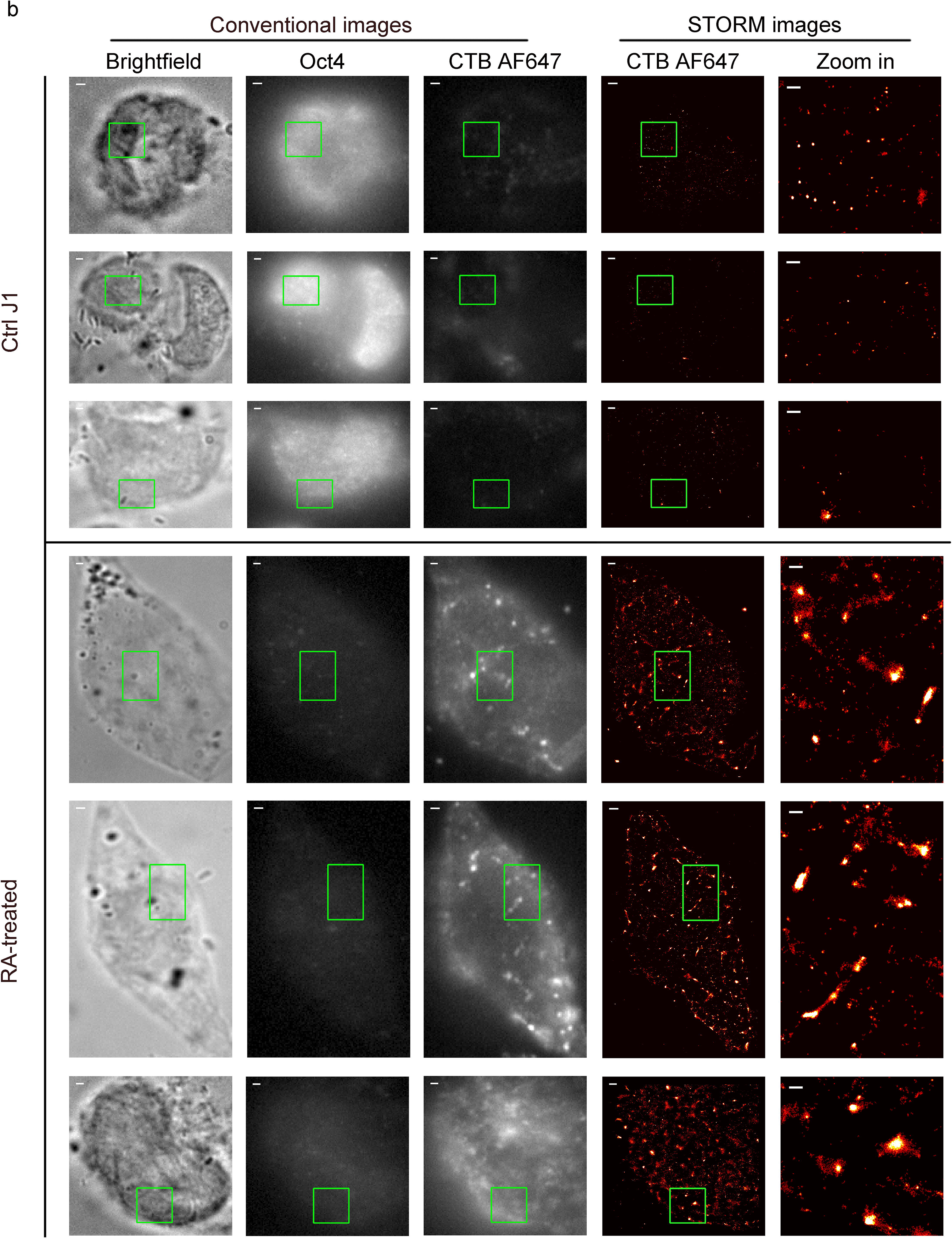
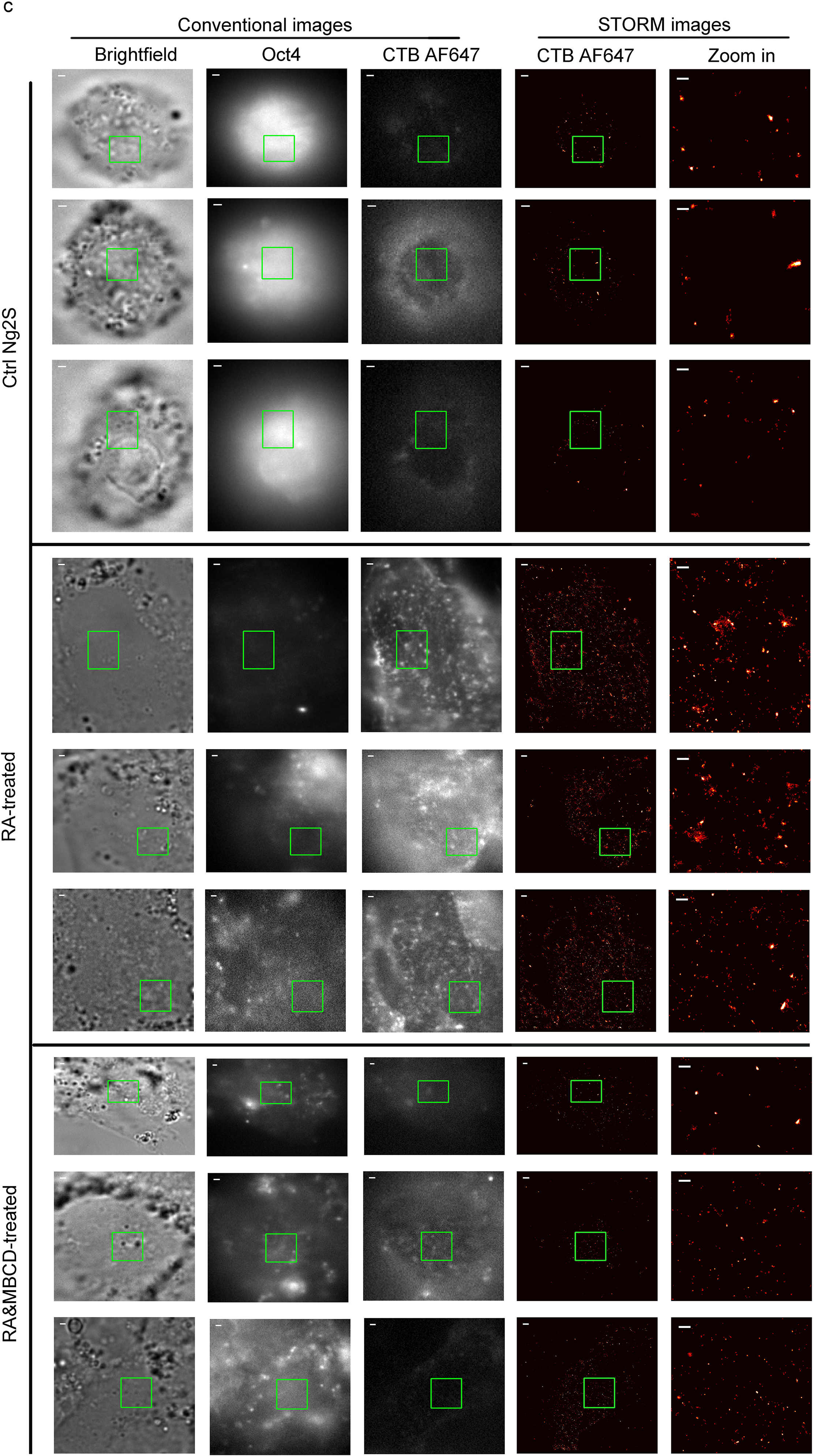
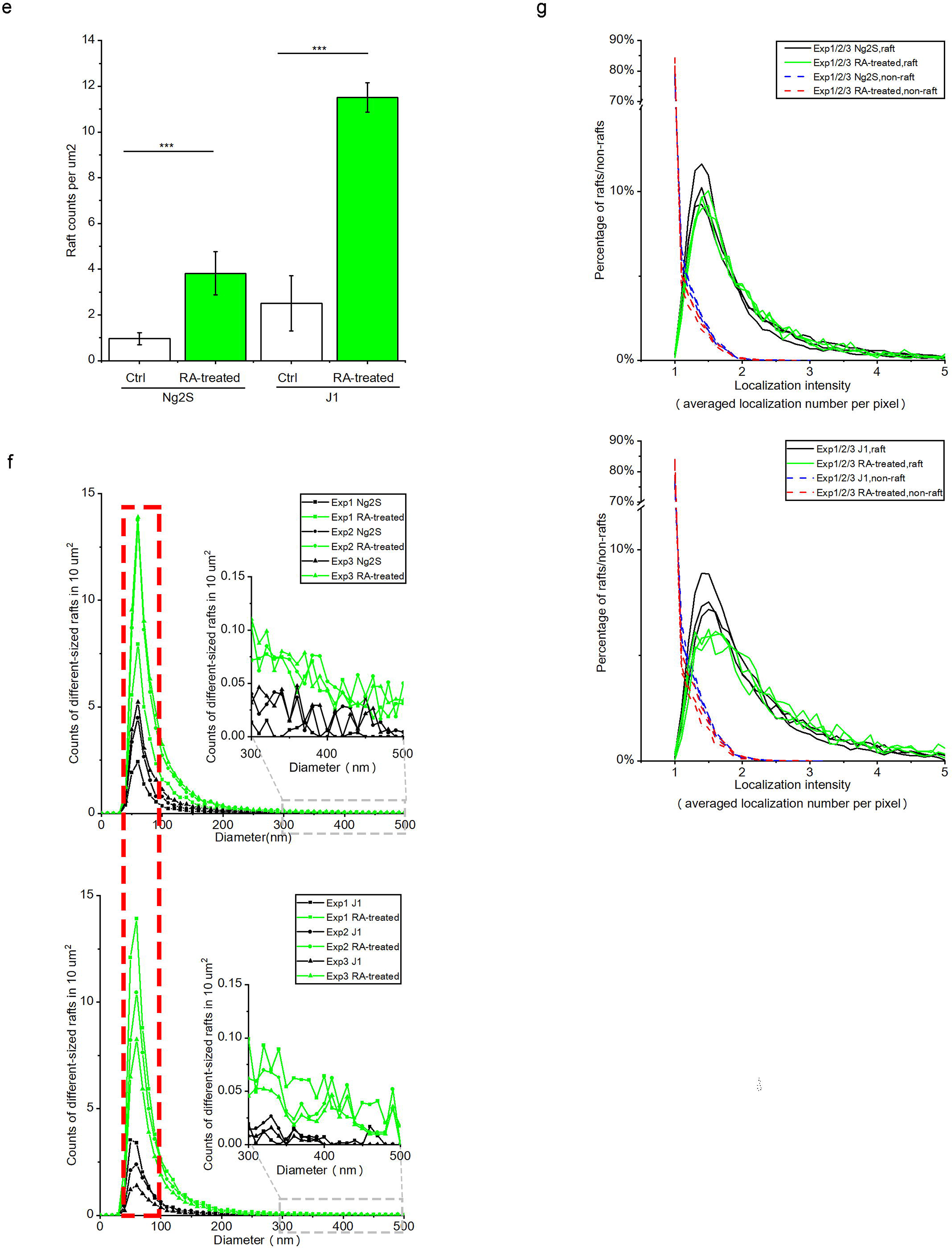
STORM characterization of distinct nanoscopic rafts in RA-treated or control mESCs. **a**, STORM imaging revealed specific raft nanostructures on J1 plasma membrane. J1 cells were labelled with CTB AF647 (upper) or control IgG AF647 (bottom) before fixation and imaging. For a certain plasma membrane region, its bright field view, conventional and STORM images of AF647 are shown in a row from left to middle. Zoom-in STORM image of the green boxed region is shown in the rightmost to reveal detailed raft structures. Scale bar for zoom-in STORM images at rightmost is 300 nm, whereas that for others is 1 μm. **b** and **c**, STORM imaging of rafts labeled by CTB AF647 in mESCs treated with or without RA. J1 (**b**) or Ng2S (**c**) was exposed to 1 μM RA or control DMSO for 96h and labeled with CTB AF647 before fixation. Oct4 was stained with specific antibody, followed by AF488 conjugated secondary antibody. For Ng2S group samples, the RA-induced differentiated cells were transiently treated with 10 uM MBCD at the end of RA exposure (**c**, bottom). For a certain plasma membrane region, its bright field view, conventional images of Oct4 and CTB AF647, and STORM image of CTF AF647 are shown in a row from left to middle. Zoom-in STORM image of the green boxed region is shown in the rightmost to reveal detailed raft structures. STORM images are presented as heat maps of single-molecular localization density: black (none) < red < yellow < white (most). Scale bar for zoom-in STORM images at rightmost is 300 nm, whereas that for others is 1 μm. **d**, Averaged counts of rafts detected in every μm^2^ area of imaged membranes of RA-treated (green bars) or control (blank bars) Ng2S or J1. The statistic results were based on STORM images of 21 pairs of RA-treated and control Ng2S and 24 pairs of RA-treated and control J1 cells from three independent experiments. Error bars, SEM. ***p<0.001; t test. **e**, The averaged counts of different sized rafts detected in every 10 μm^2^ area of imaged membranes of RA-treated (green lines) or control (black lines) cells. The parts in red box represent the most abundant subpopulation of control or differentiated Ng2S (upper) or J1 (bottom). The distribution of large rafts with diameter of 300-500 nm are zoomed as shown for a clear view. The statistic results were based on STORM images of 21 pairs of RA-treated and control Ng2S and 24 pairs of RA-treated and control J1 cells from three independent experiments. Error bars, SEM. **f**, The averaged intensity (localization number per pixel) of rafts or non-raft regions on plasma membrane of control or RA-treated cells. The percentages of rafts of different intensity are shown as black (control) and green (RA-treated) solid lines. The percentage of non-raft regions of different intensity are shown as blue (control) and red (RA-treated) dashed lines. The statistic results were based on STORM images of 21 pairs of RA-treated and control Ng2S and 24 pairs of RA-treated and control J1 cells from three independent experiments. Ctrl, control; Exp, Experiment

Next, we further assessed raft features in mESCs and their RA-induced differentiated cells at super-resolution. In conventional images (**Figure 2b**, left), Oct4 staining was remarkably weaker in RA-treated cells than control J1, confirming the RA-induced differentiation, whereas stronger CTB AF647-positive raft puncta were observed in a large number of differentiated cells rather than mESCs. But we failed to obtain further detailed features about raft puncta until applying STORM imaging. As shown in zoom-in STORM images, a dramatically more rafts of various sizes were observed distributing on plasma membrane of differentiated cells than control J1 cells (**Figure 2b**, rightmost). Very large rafts with diameters of around 300 nm were frequently observed in RA-induced cells but seldom detected in controls. Regardless of the rafts sizes, the centres of many rafts were found densely compacted according to their relatively high localization number. Consistently, STORM imaging revealed similar increase of raft abundance and sizes in many of the RA-treated cells, compared to control Ng2S, although rafts in these cells appeared a bit sparser correspondingly than those in another pair of mESC (J1) and differentiated cells (**Figure 2c**, rightmost, upper & middle). Furthermore, the increased nanoscopic structures in RA-treated cells were confirmed to be rafts, as they were significantly diminished by transient treatment with MBCD (**Figure 2c**, rightmost, bottom). These large rafts were not due to the CTB-mediated tetrameric artefact, because they were repeatedly observed in RA-treated cells that were fixed before CTB labelling (**Supplemental Figure 3**). Therefore, STORM imaging provides direct evidence at nanoscale resolution that rafts existed in mESCs and were remarkably distinct after RA-induced differentiation.

To obtain quantitative knowledge of rafts in pluripotent or differentiated cells, we analysed the raft features according to STORM images. We found that the rafts labelled with CTB AF647 covered roughly around 1.4±0.7 % or 2.2±1.2 % of the imaged plasma membrane in Ng2S or J1, respectively. After 96 h RA treatment, the coverage rate was augmented to 7.9 <u>±</u> 2.3 % or 10.6 ±7.4 %, respectively. The raft number was significantly higher in the same surface area of RA-treated cells (green bars) than control Ng2S or J1 (blank bars, **Figure 2d**). The counts of different-sized rafts revealed that the distribution density was remarkably higher in RA-induced cells (green lines) than control mESCs (black lines), while the most abundant rafts in both cells shared similar diameters peaked at 50-60 nm (red dashed rectangle, **Figure 2e**). Since rafts are relatively dense membrane structures, we then compared the packing states of rafts in different cell lines or types via assessing CTB AF647 fluorophore intensity. In our STORM images, individual clusters containing at least five localizations were regarded as rafts, whereas those containing less as non-raft regions, similar to previous reported[10]. As shown in Figure 2f, most of the identified rafts (black & green, solid lines) exhibited higher localization intensity than the non-raft regions (blue & red, dashed lines) in the same cells regardless of different cell lines or types, consisting with the concept that rafts are relatively dense compared to the surrounding membrane [1, 10]. Meanwhile, when comparing rafts in mESCs, Ng2S or J1 (black lines), to those in differentiated cells (green lines), we observed little difference in their averaged localization intensities (**Figure 2f**), indicating that raft intensity in mESCs or RA-treated cells were similar. Taken together, these results provided direct evidence at super-resolution that rafts in mESCs or their RA-induced differentiated cells are remarkably different in membrane coverage rate, distribution density, and detectability of large sizes while sharing similar most abundant population and packing state.

### Lipid rafts at early differentiation contribute to RA-induced commitment of ectoderm-like cells

Raft component sphingolipids or cholesterol has been reported to play a role in neural function and cell proliferation and differentiation, including neural differentiation from embryonic bodies or neural progenitor cells [6][39]. Much less is known about the potential relationship of rafts with early differentiation events. Glycosphingolipid depletion *in vivo* resulted in mouse embryonic lethality at E7.5, probably due to abnormality in ectodermal layer formation [6]. Therefore, we suspected that rafts are involved in lineage commitment, particularly early steps that affects cell fate determination. To address this question, we applied CRISPR/Cas9-mediated ablation of *ugcg* gene that encodes a glucosyltransferase required for glycosphingolipid synthesis [6, 40]. Then, we measured the surface level of raft component ganglioside GM1 and found remarkably lower CTB binding in five different mutant clones (orange lines) than those control clones transfected with mock vectors (grey lines, **Figure 3a**), confirming the efficient ablation of *ugcg* gene in the mutant cells. Then, we characterized RA-induced differentiated cells using antibodies against CD24 and PDGFRA after 96 h RA exposure and observed multiple cell types, including ectoderm-like (CD24+/PDGFRA-) and XEN-like (CD24-/PDGFRA+) cells. Interestingly, smaller subpopulations of ectoderm-like cells (green) were found to be derived from *ugcg* mutant clones than from the mock controls (mock group: 35.92±4.56 %, mutant group: 22.24±3.23%, **Figure 3b**). These findings showed that lipid rafts play a role in RA-induced commitment of ectoderm-like cells.

**Figure 3.**
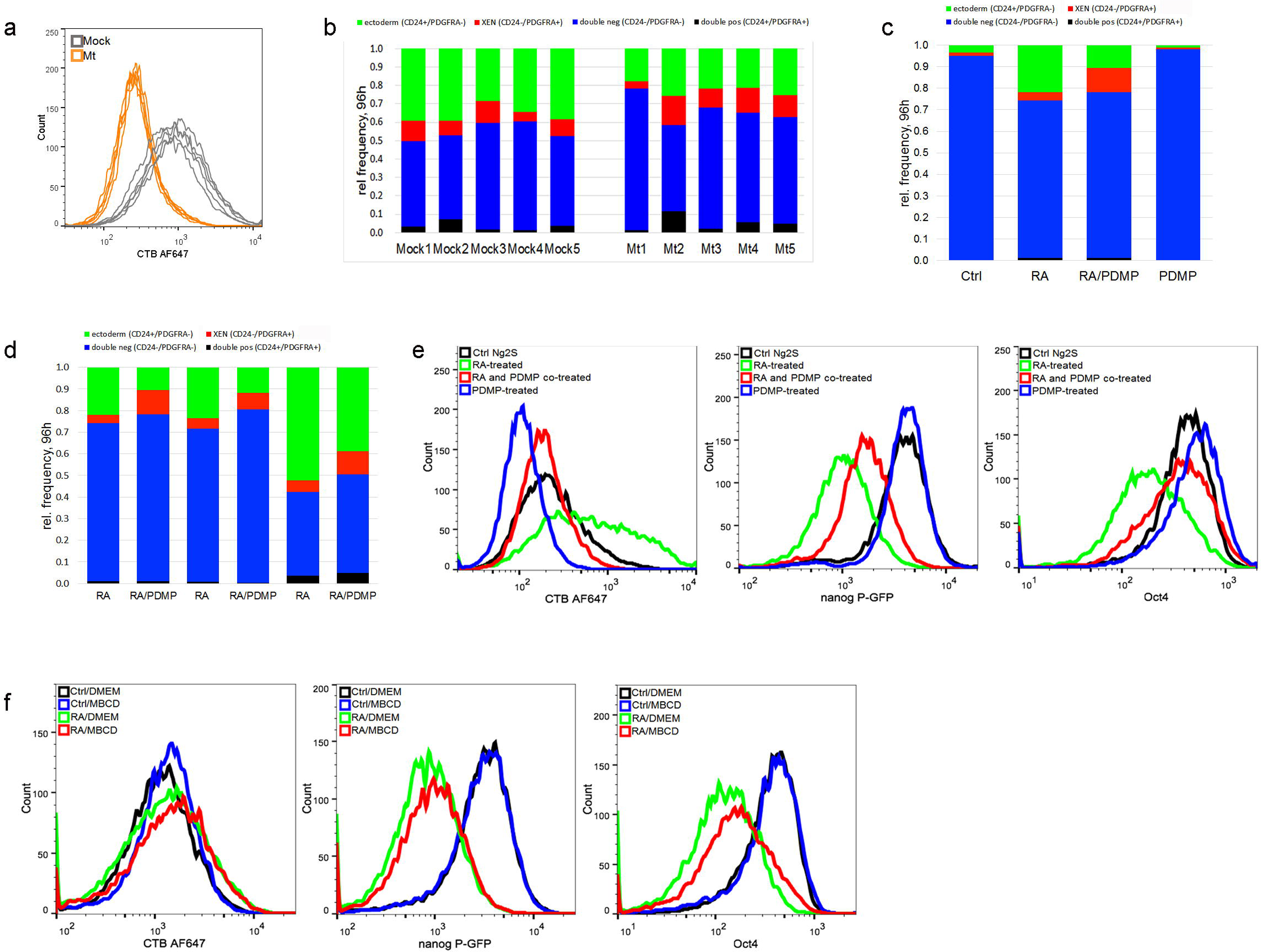
Raft inhibition diminished RA-induced commitment of ectoderm-like cells from mESCs. **a**, Frame-shift mutations of *ugcg* gene in Ng2S cells were mediated by CRISPR/Cas9. Mock controls or cells from individual clones transfected with *ugcg*-targeting gRNA were stained with CTB AF647, followed by flow cytometry quantification. Decreased CTB labeling was observed in multiple mutant clones, which representing *ugcg* ablation (Mt, orange), compared to mock control (Mock, grey). **b**, Frequency of various cell types derived from mock controls or cells with ablated *ugcg* gene after 96h exposure to RA., cells treated with RA or DMSO for 96h were subsequently co-stained with antibodies specific for CD24 and PDGFRA and measured by flow cytometry. Percentage of ectoderm-like (CD24+/PDGFRA-, green), XEN-like (PDGFRA+/CD24-, red), double negative (CD24-/PDGFRA-, blue) or double positive cells (CD24+/PDGFRA+, black) are shown in the bar graph. **c**, Frequency of various cell types derived from 96 h RA exposure. Ng2S cells were treated with DMSO (Ctrl), 1 μM RA, 30 μM PDMP or co-treated with RA and PDMP (RA/PDMP). PDMP was added 12 h preceding RA treatment and removed after 48 h RA incubation whereas RA exposure lasted continuously for 96 h. The cells were labeled with CTB AF647, co-stained with antibodies against CD24 and PDGFRA, and analyzed by flow cytometry. Percentage of ectoderm-like (CD24+/PDGFRA-, green), XEN-like (PDGFRA+/CD24-, red), double negative (CD24-/PDGFRA-, blue) or double positive cells (CD24+/PDGFRA+, black) are shown in the bar graph. **d**, Flow cytometry measurement of CD24 and PDGFRA expression for RA and RA/PDMP groups are shown from three independent experiments. **e**, Flow analysis of pluripotency gene expression of Ng2S after 48 h exposure to RA with/without PDMP treatment. Ng2S cells were treated with DMSO (Ctrl, black), 1 μM RA (green), 30 μM PDMP (blue) or co-treated with RA and PDMP (RA/PDMP, red) for 48 h. PDMP was administered for −12 h to 48 h. Cells were stained with CTB AF647 and anti-Oct4 antibodies, followed by AF568 conjugated secondary antibody. CTB labelling and expression of Oct4 and Nanog were quantified by flow analysis. **f**, Flow analysis of pluripotency gene expression of Ng2S after 48 h exposure to RA with/without MBCD treatment. Ng2S cells were treated with 1 μM RA or DMSO (Ctrl) for 48 h, during which 10 mM MBCD or DMEM was administered to cells for 50 min at −1 h and 12 h. Cells stained with CTB AF647 and anti-Oct4 antibodies were analyzed with flow cytometry. Representative results are shown from three independent experiments.

In response to RA treatment, raft increase appeared from the 12 h time point and reached a profound level before pluripotency exit. These findings implied a potential role of rafts in early differentiation, which has been rarely reported. To address this question, we set out to inhibit rafts at the early stage of RA-induced differentiation with the use of glycosphingolipid synthase inhibitor PDMP (D-threo-l-phenyl-2-decanoylamino-3-morpholino-l-propanol) [26, 41, 42]. As expected, 96 h RA exposure of Ng2S resulted in the differentiation of a substantial amount of ectoderm-like cells (CD24+/PDGFRA-, green), whereas co-treatment with PDMP for the first 36-48 h reduced the subpopulations (**Figure 3c**). Nonetheless, the differentiation state was similar in both cases after RA exposure, as indicated by the GFP expression (**Supplemental Figure 4a**). At the end, the raft level in cells undergoing early stage raft inhibition was still observed to be slightly lower than that in the controls (**Supplemental Figure 4b**). The inhibitory effects of raft disruption on RA-induced ectoderm-like cell commitment was relatively robust across multiple independent experiments (**Figure 3d**). Therefore, these results showed that rafts in the early differentiating cells play a role in facilitating ectoderm-like cell commitment during RA exposure.

To our current knowledge, expression of pluripotency factors, such as Oct4 and Nanog, was reduced in differentiating mESCs, representing pluripotency exit before undergoing lineage specification [43-46]. We tested if lipid rafts also contribute to pluripotency exit by treating Ng2S with RA alone or RA together with PDMP. As expected, expression of Oct4 and GFP reporter decreased while rafts increased in RA-treated cells (green lines), compared to control mESCs (black lines, **Figure 3e**). Interestingly, co-treatment with PDMP (red lines) weakened the RA-induced effects (green lines), retarding the raft increase and the pluripotency marker decrease (**Figure 3e**). These results indicated that lipid rafts play a role in RA-induced exit from pluripotency. To verify this indication, we also disrupted rafts via treating the early differentiating cells with cholesterol depletion reagent MBCD [47-50]. Compared to the group treated with RA only (green), cells co-treated with MBCD and RA exhibited delayed pluripotency exit, as revealed by the slower decrease of Oct4 and GFP expression (red, **Figure 3f**), supporting that rafts facilitate the early differentiating cells to exit from pluripotency. Taken together, these data showed that lipid rafts play a role in facilitating ectoderm-like cell commitment as well as exit from pluripotency during RA-induced differentiation.

## Discussion

This study demonstrated that cell-surface lipid rafts remarkably increase when the monolayer-cultured mESCs (J1, Ng2S, and Cj9) were committed into differentiated cells, particular ectoderm-like cells, in response to RA exposure. STORM imaging demonstrated discrete nanoscopic rafts on plasma membrane of mESC and a remarkable increase in raft distribution density in RA-induced differentiated cells. For the first time, this study demonstrated at super-resolution the existence and distinct features of cell-surface rafts in mESCs and there differentiated cells, although there are still controversial views on the existence of rafts in cells. CRISPR-mediated knockout of *ugcg* resulted in remarkable depletion of cell-surface rafts and decreased RA-induced subpopulation of ectoderm-like cells. These results showed lipid rafts are involved in ectoderm-like cell differentiation, which possibly contribute to ectodermal layer abnormality in *ugcg* knockout mice [6]. Furthermore, raft inhibition in early differentiating stage of RA-induced cell commitment had a similar inhibitory effect on ectoderm-like cell subpopulation, indicating that rafts in the early differentiating stage contribute to cell fate determination. Meanwhile, we observed rafts also contributed to exit of pluripotency that is governed by a transcription factor internet. Therefore, lipid rafts play a role in the early differentiating stage, facilitating pluripotency exit and ectoderm-like cell determination.

## Materials and Methods

### Materials

All-trans retinoic acid (RA, R2625, Sigma-Aldrich) was dissolved in dimethyl sulfoxide (DMSO) to the stock concentration of 10 mM. D-threo-1-phenyl-2-decanoylamino-3-morpholino-1-propanol (D-threo-PDMP, shorted as PDMP, #1756, Matreya LLC) was dissolved in DMSO at the stock concentration of 300 mM. Methyl-β-cyclodextrin (MBCD, C4555, Sigma-Aldrich) dissolved in DMEM. Alexa Fluor 647-Conjugated Cholera Toxin Subunit B (CTB AF647, C34778, Invitrogen) was dissolved in PBS at the stock concentration of 1 mg/mL. Filipin III from Streptomyces filipinensis (Filipin, F4767, Sigma-Aldrich) was dissolved in DMSO at the stock concentration of 25 mg/mL.

### Mouse embryonic stem cell (mESC) culture and treatments

mESC J1 was purchased from ATCC. Ng2S is generated from a mESC line CCE through introducing GFP gene under the nanog promoter. mESC Cj9 was kindly provided by Dr. Xiaohua Shen (Tsinghua University, China). All mESC lines were cultured in feeder free condition in complete ESC culture medium containing DMEM (Dulbecco’s modified Eagle’s medium, 10-013-CVR, Corning) supplemented with 15% fetal bovine serum (FBS, 10099-141, Gibco), 0.1 mM MEM Non-Essential Amino Acids (11140050, Gibco), 2 mM Glutamax (35050061, Gibco), 0.1 mM 2-Mercaptoethanol (AMRESCO), 1000 U/mL mouse recombinant Leukemia inhibitory factor (LIF, ESG1107, Millipore) and Penicillin-Streptomycin-Neomycin Antibiotic Mixture (15640055, Gibco). Cells were passaged every other day with Accutase (A1110501, Gibco) and replated on gelatinized tissue culture plates. Culture medium was refreshed every day.

The mESCs were grown around 2 passages after revival and then subjected to various assays. RA treatment was carried out continuously in ESC culture medium without LIF supplemented with 1 μM RA. Spent medium was exchanged with fresh medium every day. For PDMP treatment, 30 μM PDMP was added 12 h prior to RA or control exposure and continuously present in culture medium for the indicated time. For MBCD treatment, cells were transiently treated with -MBCD of the indicated concentration at 37°C for 40 min at the indicated times.

### CTB AF647 staining, immuno-fluorescence staining and microscopy

Cells on coverslips (12-545-82, Fisher, Pittsburgh, PA) were cultured for the indicated times. For CTB staining, unless otherwise mentioned, live cells were stained with CTB AF647 in DMEM at the concentration of 2 μg/mL at 18°C for 10 min and fixed with 4% PFA at room temperature for 10 min. The fixed cells were permeabilized with 0.5% Triton-X-100 in PBS for 10 min and blocked in blocking buffer consisting 5% donkey serum and 2% BSA in PBS for 30 min at room temperature. For Oct4 staining, cells were incubated with antibody against Oct4 (sc-5279, Santa Cruz) at 1:200 dilution at 4D overnight, followed by 1h incubation with AF488-conjugated secondary antibody (A21202, Invitrogen). Cell samples were mounted in SlowFade Gold Antifade Mountant with DAPI (S36938, Invitrogen) for imaging using an inverted wide-field fluorescent microscope (Ti, Nikon) with a 20x dry objective. The wide-field images were captured by camera (dmk23u445, Imaging Sourse) in their corresponding fluorescence band-pass filters combinations: Semrock 49000 for dapi, 49011 for AF 488,49008 for AF 568 and 49006 for AF 647.

### Flow analysis

For live cells stained with 2 μg/mL CTB AF647 in DMEM were washed with PBS once, dissociated with Accutase, and collected by centrifugation at 400 g for 2 min before fixation with 4% Paraformaldehyde (PFA, Sigma-Aldrich) at room temperature for 10 min. For intracellular protein staining, cells were permeabilized with 0.5% Triton-X-100 in PBS for 10 min. The samples were blocked in blocking buffer consisting 5% donkey serum and 2% BSA in PBS for 30 min at room temperature, before incubating with antibodies for 1 h at room temperature. Anti-Oct4 antibody (ab19857, abcam) and anti-Nanog antibody (ab80892, abcam) were used at 1: 1000 and 1: 200 dilution in blocking buffer, respectively, followed by 1 h incubation of AF-568- or AF647-cojugated secondary antibodies (A21206 and A10042, Invitrogen). For staining of membrane protein CD24 and PDGFRA (CD140a), PE rat anti-mouse CD24 antibody (553262, BD Pharmingen) and Super Bright 436-conjugated anti-CD140a (PDGFRA) antibody (62-1401-80, Invitrogen) were used at 1:1000 and 1:250 dilution. To measure the amount of cholesterol, fixed cells were stained with filipin at the concentration of 0.125 mg/mL in PBS for 20 min at RT. Cells samples were analyzed using BD FACS cytometer.

### CRISPR/Cas9-mediated knockout of *ugcg* gene

Single guide RNAs (sgRNA) targeting *ugcg* gene were designed using the CRISPR design tool (http://crispr.mit.edu/). The targeting sequences of sgRNAs (PAM sites are underlined) were #1, GTTCGGCTTCGTGCTCTTCG**<u>TGG</u>** and #2, CAGGAGGGAATGGCCTTGTT**<u>CGG</u>**.

The sgRNA sequences were cloned into the sgRNA expression vector pSpCas9 (BB)-2A-Puro (PX459). 1mg sgRNA plasmid was transfected into 9×10^4^ Ng2S cells by Lipofectamine 2000 Transfection Reagent (11668019, Invitrogen). Culture medium was refreshed after 12 h to remove Lipofectamine 2000 Transfection Reagent. At 48 h post transfection, transfectants were selected with 3.0 μg/mL puromycin dihydrochloride (A1113803, Gibco) for 36h and then plated onto 6 cm dishes at clonal density. After another 4 days, individual clones (20-24 clones) were picked and expanded for 4 days.

In order to verify the effect of frame-shift mutations, cells were stained with CTB AF647 and quantified by flow cytometry. Multiple mutant clones with *ugcg* ablation and mock controls were selected and treated with DMSO or RA for further study.

### STORM imaging

A STORM system based on an inverted optical microscope (IX-71, Olympus) with a 100x oil immersion objective lens (UplanSApo, N.A. 1.40, Olympus) was used for the nanoimaging as previously described [38]. A 405 nm laser (CUBE 405–100C; Coherent) was used for photoactivation and a 641 nm laser (CUBE 640–100C; Coherent) was used to excite fluorescence and deactivate Alexa-647 to the dark state. A 488nm laser(Sapphire 488; Coherent) was used for conventional imaging of the labelled-Oct4. The highly inclined and laminated optical sheet (HILO) configuration [51] was adopted for illumination to diminish/minimize defocused background. A dichroic mirror (Di03-r405/488/561/635 t1, Chroma) was used to separate the fluorescence from the laser and a band-pass filter (FF01-676/37 or FF01-514/30, Semrock for AF647 or AF488, Semrock) on the imaging path was used to filter the fluorescence. Fluorescent signals of each cell-surface were acquired with an EMCCD (DU-897U-CV0, Andor) at 46fps unless otherwise indicated. The 641 nm laser was kept on during the whole imaging process, and the 405 nm laser was added only in the second series of images when necessary. An auto-correlative algorithm was applied to correct for drift correction. Unless specified, a standard STORM imaging buffer containing 50 mM Tris (pH 8.0), 10 mM NaCl, 1% b-mercaptoethanol (v/v), 10% glucose (w/v), 0.5 mg/mL glucose oxidase (G2133, Sigma), and 40 m g/mL catalase (C30, Sigma) [52, 53]was used.

### STORM image reconstruction and raft identification

The STORM imaging data were analyzed in a similar manner as previously described [38]. Briefly, the fluorescence images of each fluorophore were preprocessed using a band-pass filter to reduce the background noise, and then fitted to a Gaussian function where the position of the fluorophore in the x-y plane was directly read out. In the reconstructed image, each localization was represented as a spot whose diameter matched with the localization precision, i.e. 22 nm [38]. Only those clusters containing >5 localizations were determined as raft[10] then painted in the super-resolution images and subjected to further analysis. The sizes of variously shaped clusters were determined with the diameters of the transformed circular clusters whose areas were equal to those original ones. Cluster intensity was determined as average gray of the region in the super-resolution images.

## Acknowledgement

We thank TS Ma, W Jin, and XH Shen for advice on mESC culture and Cj9, RL Sun and LZ Liu for supports in CRISPR/Cas9 knockout, Bin Yu and YJ Sun for support and constructive suggestions on imaging, N Zhou, QB Lin, L Rong, BZ Zhang, J Deng, SW Li, Teng Luo, BL Chen, Y He, X Peng, and J Man for their technical assistance. This work was supported by the National Natural Science Foundation of China (31401146, 31871293, 61235012), the Special-Funded Program on National Key Scientific Instruments and Equipment Development (2012YQ150092), the National Basic Research Program of China (2012CB825802, 2015CB352005, 2012CB316503), and the Shenzhen Science and Technology Planning Project (JCYJ20170817095211560, JCY20170817035211560).

## Figure legends

**Supplement Figure 1** Confocal images of GM1 labeling at different temperature by CTB AF647 on J1. J1 cells treated with RA or DMSO (Ctrl) for 96h were labelled by CTB AF647 at 12°C, 18°C, and 24°C. Confocal images were captured using a Leica SP2 confocal microscope using a 63x oil objective and 633nm laser. The confocal microscopy images are shown in column 1-3, respectively. The views of bright field are shown in row 1 and 4, the corresponding fluorescence images are shown in row 2 and 3. Scale bar is 10 μm.

**Supplemental Figure 2** Cj9 cells on coverslips were cultured with 1 μM RA or DMSO (Ctrl) for 96 h, labelled with CTB AF647. The fixed cells were stained with anti-Oct4 antibody, followed by AF488 conjugated secondary antibody. The coverslips were mounted using SlowFade Gold Antifade Mountant with DAPI. Samples were captured by Nikon-Ti using 405 nm, 488 nm and 641 nm lasers, or under bright field. Scale bars are 10 μm.

**Supplement Figure 3** STORM images of lipid rafts labeled by CTB AF647 after fixation in Ng2S cells treated with RA or control DMSO for 96h. Oct4 was stained with specific antibody, followed by AF488 conjugated secondary antibody. For a certain plasma membrane region, its brightfield view, conventional images of Oct4 and CTB AF647, and STORM image of CTB AF647 are shown in a row from left to middle. Zoom-in STORM image corresponding to the green boxed region is shown in the rightmost to reveal detailed raft structures. STORM images are presented as heat maps of single-molecular localization density: black (none) < red < yellow < white (most). Scale bar for zoom-in STORM images at rightmost is 300 nm, whereas that for others is 1 μm.

**Supplement Figure 4** Flow analysis of CTB labeling and Nanog expression of Ng2S after 96 h exposure to RA with/without PDMP treatment. Ng2S cells were treated with DMSO (Ctrl, black), 1 μM RA (green), 30 μM PDMP (blue) or co-treated with RA and PDMP (red) for 96 h. PDMP was administered from −12 h to 48 h. After 96 h, cells were stained with CTB AF647, anti-CD24 and anti-PDGFRA antibodies. Expression of CD24 and PDGFRA are displayed in Figure 3c.

