## Supplementary figures and images for "Lipid rafts increase to facilitate ectoderm lineage specification of differentiating embryonic stem cells"

### Supplemental Figure 1

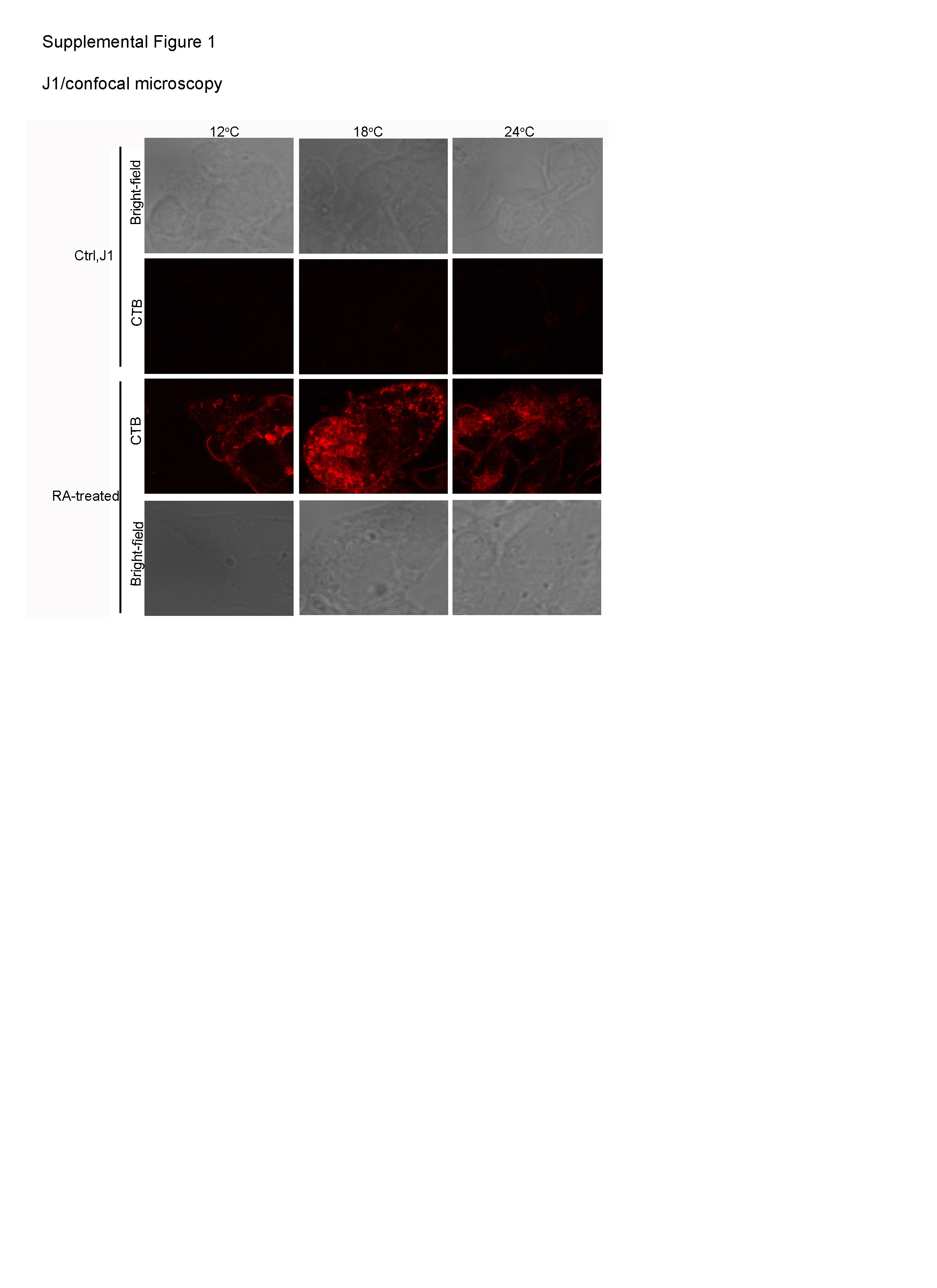

### Supplemental Figure 2

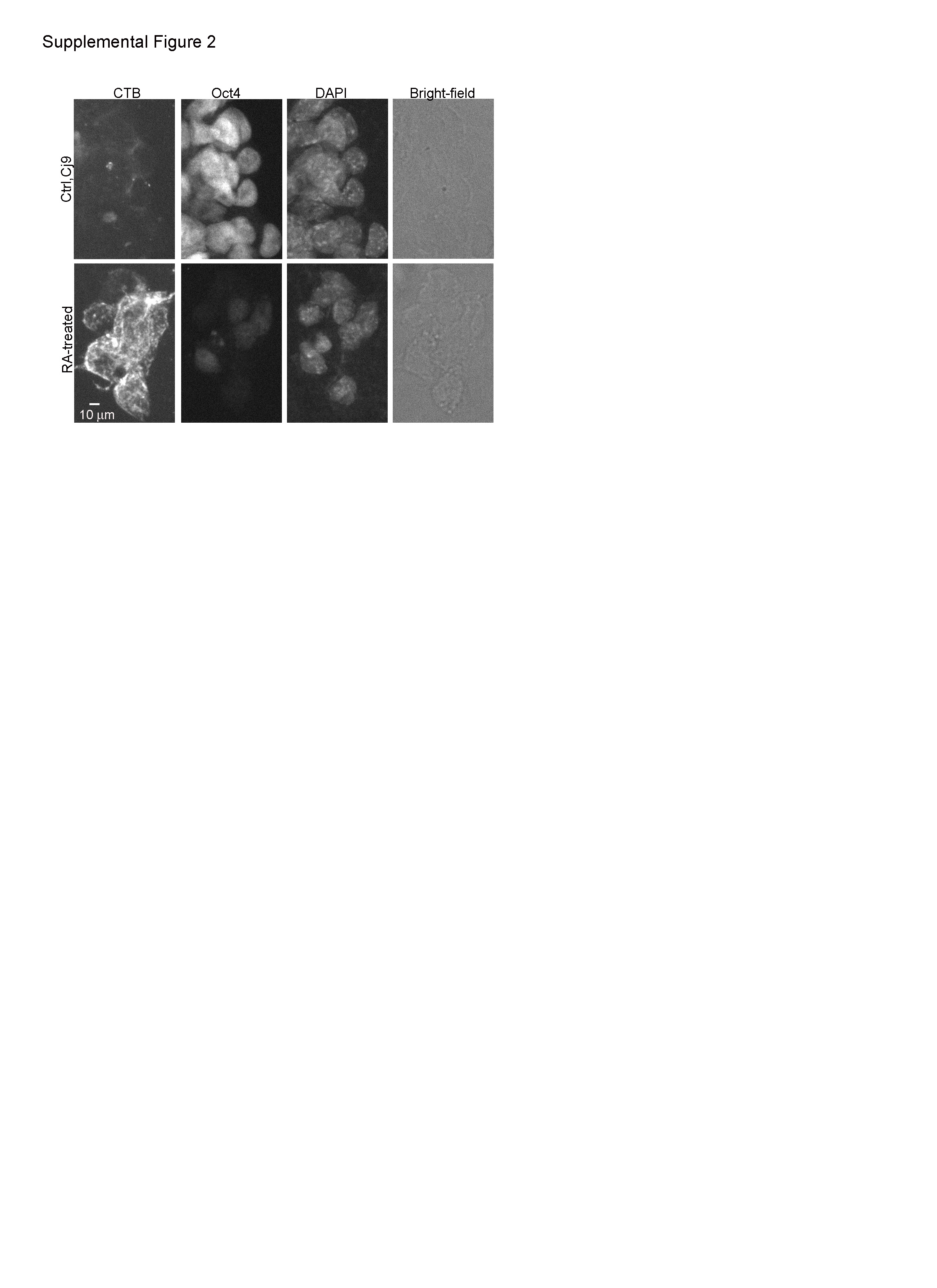

### Supplemental Figure 3

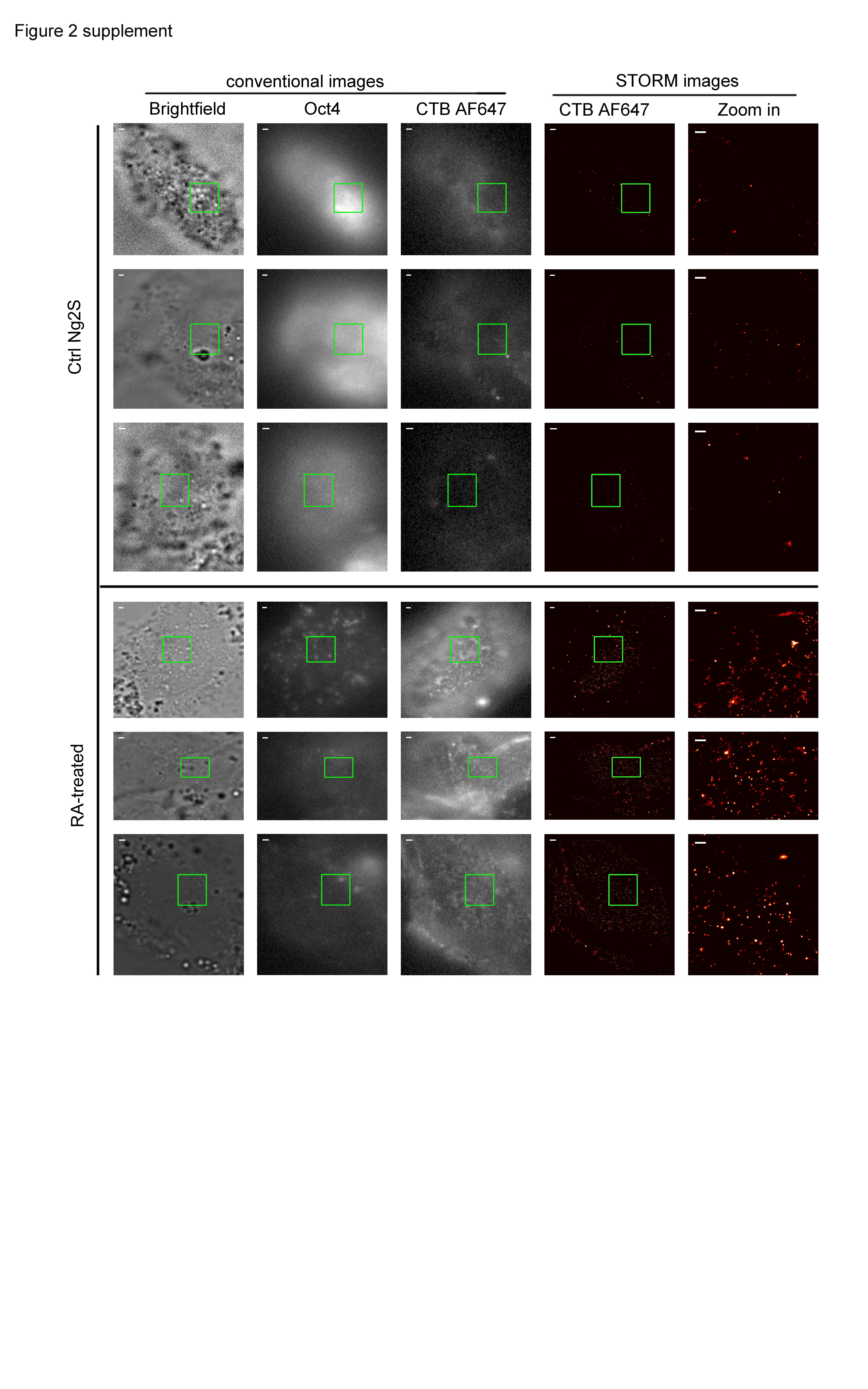

### Supplemental Figure 4

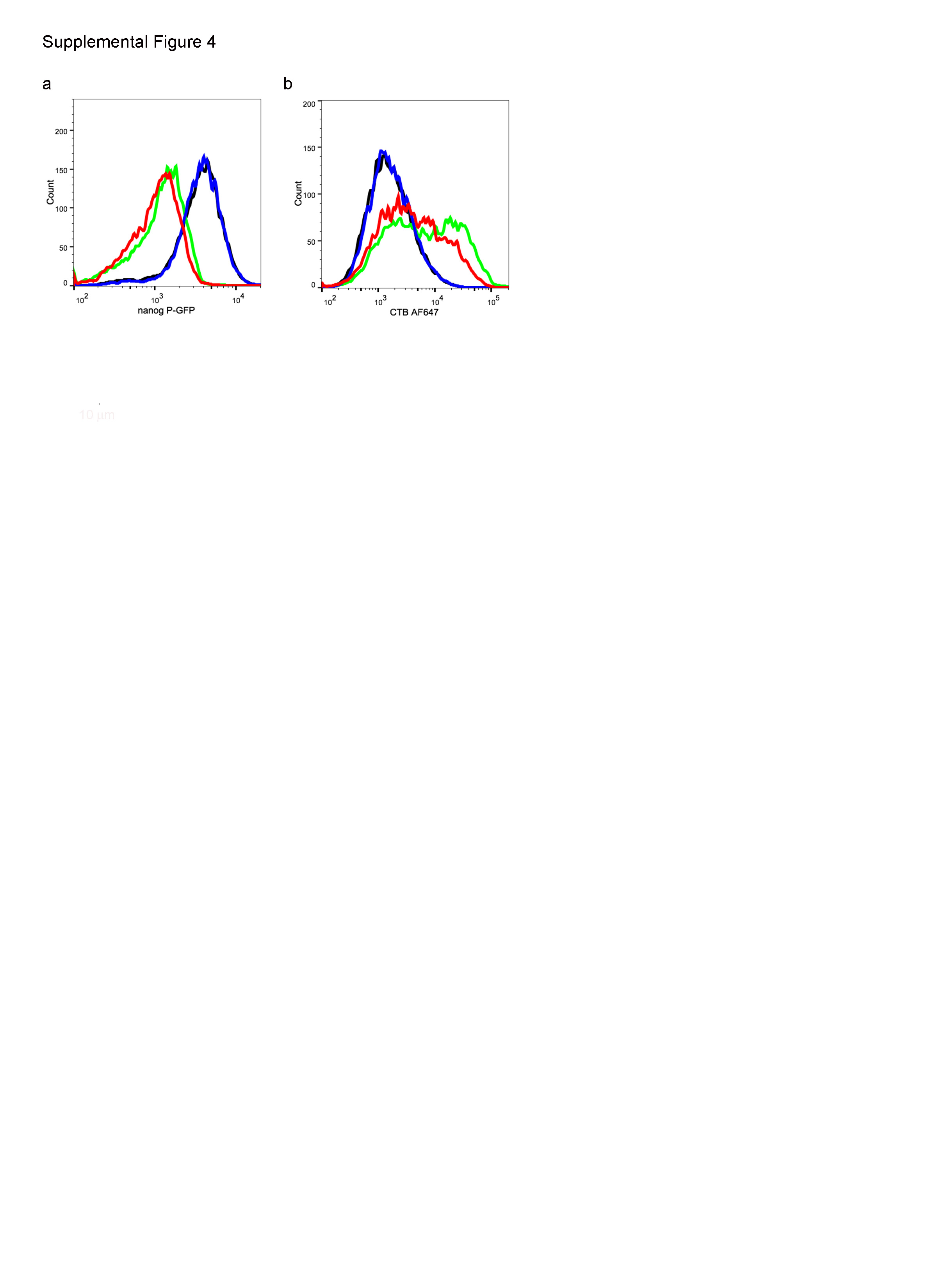
